# A metabolomics insight into the Cyclic Nucleotide Monophosphate signaling cascade in tomato under non-stress and salinity conditions

**DOI:** 10.1101/2021.03.22.436432

**Authors:** Begoña Miras-Moreno, Leilei Zhang, Biancamaria Senizza, Luigi Lucini

## Abstract

Cyclic Nucleotides Monophosphate (cNMP) are key signalling compounds whose role in plant cell signal transduction is till poorly understood. In this work we used sildenafil, a phosphodiesterase (PDE) inhibitor used in human, to amplify the signal cascade triggered by cNMP using tomato as model plant. Metabolomics was then used, together with plant growth and root architecture parameters, to unravel the changes elicited by PDE inhibition either under non-stress and 100 mM NaCl salinity conditions.

The PDE inhibitor elicited a significant increase in biomass (+62%) and root length (+56%) under no stress conditions, and affected root architecture in terms of distribution over diameter classes. Together with cGMP, others cNMP were modulated by the treatment. Moreover, PDE inhibition triggered a broad metabolic reprogramming involving photosynthesis and secondary metabolism. A complex crosstalk network of phytohormones and other signalling compounds could be observed in treated plants. Nonetheless, metabolites related to redox imbalance processes and NO signalling could be highlighted in tomato following PDE application. Despite salinity damped down the growth-promoting effects of sildenafil, interesting implications in plant mitigation to stress-related detrimental effects could be observed.

**HIGHLIGHT:** The role of Cyclic Nucleotides Monophosphate in plant cell signal transduction involves regulation of plant growth and architecture, together with a broad biochemical reprogramming of metabolism.

## 1. INTRODUCTION

In the last years, an increasing interest in cyclic nucleotide monophosphates (cNMP - in particular, in cAMP and cGMP) has grown, driven by the pivotal role they might play in plant growth and in response to environment. The role of cNMP in plants has been argued for long time, because they are present in nanomolar concentrations, i.e. one order of magnitude lower than in mammalian cells. Furthermore, a set of plant cyclases has been proposed, including membrane protein and cytosolic enzyme, with >50 candidate proposed in the *Arabidopsis thaliana* proteome having different different domain organizations (Wong and Gehring, 2013). Both adenylate and guanylate cyclases have been defined “moonlighting” proteins, since they are a single polypeptide chain holding diverse domain structures and harbouring multiple catalytic domains (Blanco *et al*., 2020). These cyclases determine the endogenous synthesis of cNMP whereas phosphodiesterase (PDE) activity determines their rate of degradation by cleaving the phosphodiester bond to yield the inactive non cyclic NMP (Swiezawska *et al*., 2018). In this way, PDE regulate the amplitude of cNMP levels and the duration of their signal in the cell (Duszyn *et al*., 2019). Although the most of scientific evidence in plant physiology relates to cGMP and cAMP, other so-called “non canonical” cNMP such as cytidine, inosine, uridine, and 2′-deoxythymidine 3′,5′-cyclic monophosphate (cCMP, cIMP, cUMP, and cdTMP, respectively) are emerging(Gehring and Turek, 2017).

Cyclic NMP are key signal transduction compounds, able to directly activate cyclic nucleotide gated channels (CNGCs) rather than cNMP dependent kinases like cAMP (PKA) and cGMP (PKG) dependent kinases, thus leading to the phosphorylation of cGMP and cAMP responsive transcription factors (Martinez-Atienza *et al*., 2007). Recent evidence indicates that cNMP interactome has binding candidates displaying critical functions in the Calvin–Benson cycle, in photorespiration (Donaldson *et al*., 2016), cell division, growth and differentiation, germination, and and flowering (Gehring and Turek, 2017). Furthermore, a complex interplay among cNMP and different signalling compounds has been proposed, including an antagonistic effect with abscisic acid and interaction with Ca^2+^ current (Yuan *et al*., 2017). The involvement of cGMP in germination and its modulation of gibberellins-mediated protein kinases suggests its tight relationship also with gibberellins (Shen *et al*., 2019). Moreover, the cGMP mediated dampening of glycolate oxidase in response to pathogens suggested it is implicated in the cross talk between NO and H_2_O_2_ signaling during hypersensitive responses (Irving *et al*., 2018). Similarly, cNMP have been linked to abiotic stress response in plant, via the regulation of CNGCs ion channels together with calmodulin (Arazi *et al*., 2000) and via their involvement in the NO signalling cascade together with salicylic acid (Gehring and Turek, 2017).

Despite being well recognised as fundamental signals in plant, the understanding about their functions in plant, their cooperation with other messengers and their transduction into cellular and physiological responses is still limited (Isner and Maathuis, 2018). This is even more evident for non-canonical cNMP (Marondedze *et al*., 2017). Considering that cNMP are expected to act via spatial and temporal clustering resulting from the stimulus-dependent triggering of cyclases, and assuming their time-transient nature, it becomes complicated to study how intracellular cNMP are perceived and transduced in plant. The use of the so-called “cAMP-sponge”, *i*.*e*. compounds able to specifically bind cAMP in vitro, is an elegant approach but it is insensitive to other cNMP (Blanco *et al*., 2020). However, the use of PDE inhibitors has been suggested as a solution to highlight cNMP role in plant. Sildenafil citrate (SC), a PDE inhibiting drug used in humans to threat erectile disfunction, has shown the ability to modulate NO-triggered cGMP levels in plant in a dose-dependent manner (Salmi *et al*., 2007). Consistently, SC was able to promote adventitious rooting in auxin-depleted explants (Pagnussat *et al*., 2003).

On these bases, our work aimed to point out SC-mediated cNMP role in plant, using a metabolomics approach to gain a holistic view into their downstream signaling components and contributing to unveil cNMP role in plant. To this object, tomato has been chosen as model plant since it is a fruit bearing crop with a worldwide distribution, commercial importance and complex secondary metabolism (Kimura and Sinha, 2008; Ganugi *et al*., 2020). Furthermore, the SC-mediated cNMP modulation of plant metabolism was investigated either under non-stress conditions or under salinity, in order to shed light also on the specific contribution of cNMP in tomato response to abiotic stress.

## 2. MATERIALS AND METHODS

### 2.1. Growth conditions, plant material and experimental design

The experiment was conducted in June 2020 at the Università Cattolica del Sacro Cuore (Piacenza, Italy) and was designed to elucidate the effect of the phosphodiesterase inhibitor on plants under both saline stress and non salt-conditions. Seedlings of tomato (cv. Heinz 3402) at the four-true-leaf stage were transplanted into a jar containing commercial soil. Composition of soil (Compo Bio) used: neutral sphagnum peat, green composted soil conditions and sand. pH in water 7.0, electrical conductibility 0.6 ds/m, density 375 kg/m3 and porosity 80% v/v.

Each experimental unit consisted in 10 plants. For the saline stress treatment, a 100 mM NaCl solution was provided in alternate days starting at Day 0. Distillate water was used for control plant and SC-treated plants. 500 μM SC (Merck) was foliarly applied every day. When NaCl was added, SC was applied 3 hours prior to the stress induction. Samples were collected at three different time-point (1, 2 and 7 days) considering five replicates per metabolomic analysis.

Entire root systems were scanned using an Epson Perfection V700 Photo scanner; the images were processed to determine the distribution of root length over diameter classes (0 - 0.1 mm; 0.1 - 0.2 mm; 0.2 - 0.3 mm; 0.3 - 0.4 mm; 0.4 - 0.5 mm; 0.5 - 0.6 mm; 0.6 - 0.8 mm; 0.8 - 1.0 mm; >1.0 cm) using WinRHIZO (Regent Instrument Inc., Canada).

### 2.2. Untargeted metabolomics

Tomato leaves (1.0 g) were extracted by homogenizer-assisted solvent extraction (Ultra-Turrax; Polytron PT, Switzerland) in 10 mL of a methanol/water solution (80:20, v/v), centrifuged and filtered through a 0.22 µm membrane as previously described (Paul *et al*., 2019). For the untargeted metabolomics, an Agilent 6550 iFunnel quadrupole-time-of-flight mass spectrometer and an Agilent 1200 series ultra-high-pressure liquid chromatographic system (UHPLC-ESI/QTOF-MS) were employed as previously reported (Senizza *et al*., 2019). Briefly, reverse phase chromatographic separation was achieved under a water-acetonitrile gradient elution, starting from 6% acetonitrile to 94% in 33□min. The mass spectrometer worked in SCAN mode with range from 100 to 1200□m/z, positive and negative polarity and extended dynamic range mode (nominal mass resolution = 30,000 FWHM). The injection sequence was randomized, injection volume was 6□μL, and six independent biological replicates were analyzed per treatment. Quality Controls (QCs) were made by pooling all the extracts and acquired throughout the sequence of analysis. QCs underwent the same chromatographic conditions used for samples but acquired in data-dependent MS/MS mode (1 Hz, 50–1200 m/z, 12 precursors per cycle), at different collision energies (10, 20 and 40 eV) as previously reported (Miras-Moreno *et al*., 2020).

The raw mass features obtained from UHPLC-ESI/QTOF-MS were processed by using the Agilent Profinder B.06 (Agilent Technologies) software. In particular, a 5-ppm tolerance for mass accuracy, following mass and retention time (0.05 min as maximum shift) alignments were used as post-acquisition processing. The database PlantCyc 12.6 (Plant Metabolic Network, http://www.plantcyc.org) was used for the annotation of features, by combining monoisotopic mass, isotopes ratio and isotopes spacing. Thereafter, MS/MS confirmations from QCs was carried out using the software MS-DIAL 4.24 (Tsugawa *et al*., 2015). To this aim, publicly available MS/MS experimental spectra built in the software (Mass Bank of North America) as well as MS-Finder *in-silico* fragmentation (using a fragmentation score > 5) against the PlantCyc database (Tsugawa *et al*., 2016) were used. The annotation process corresponded to level 2 of confidence as set out in COSMOS metabolomics standards initiative (Salek *et al*., 2015).

### 2.3. Statistical analysis

Results on growth parameters were expressed as means with standard deviation, and one-way analysis of variance (ANOVA) followed by Dunnett’s multiple comparisons tests was performed using Prism GraphPad software, Version 6.01 (GraphPad Software, Inc.). Elaboration of metabolomics data was carried out by using the software Mass Profiler Professional 12.6 (Agilent Technologies) for log2 transformation, normalization at the 75^th^ percentile, and baselining against the median of each compound. Thereafter, both unsupervised and supervised multivariate statistics were applied for interpretations. The unsupervised hierarchical cluster analysis was used to underline the relatedness across the different treatments, according to Euclidean distance and Ward’s linkage. In addition, the orthogonal projection to latent structures discriminant analysis (OPLS-DA) was carried out as supervised tool, recording goodness-of-fit R2Y and goodness-of-prediction Q2Y. Then, the Variable Importance in Projection (VIP) selection method was used to select the metabolites having the highest discriminant potential (VIP score > 1.20). Finally, a volcano plot analysis was applied to identify differential compounds by combining ANOVA (p < 0.05; Bonferroni multiple testing correction) and fold-change analysis (cut-off ≥ 2). Finally, differential compounds were exported into the Omic Viewer Pathway Tool of PlantCyc (Stanford, CA, USA) software (Caspi *et al*., 2013) for interpretation.

## 3. RESULTS

### 3.1 Effect of the inhibitor of phosphodiesterases on plant biomass and root architecture

Firstly, the effect of the inhibitor of phosphodiesterases (SC) was evaluated in terms of plant biomass and root architecture. The biomass was strongly affected by SC in non-stressed plants, throughout the 7 days of treatment (Figure 1A). In more detail, after 7 days of treatment SC-treated plants presented an increase of FW of +62% compared to the control (Figure 1B). However, although an obvious decrease in biomass could be observed under salinity, the treatment with SC did not induce differences in biomass in the presence of 100 mM NaCl.

**Figure 1.**
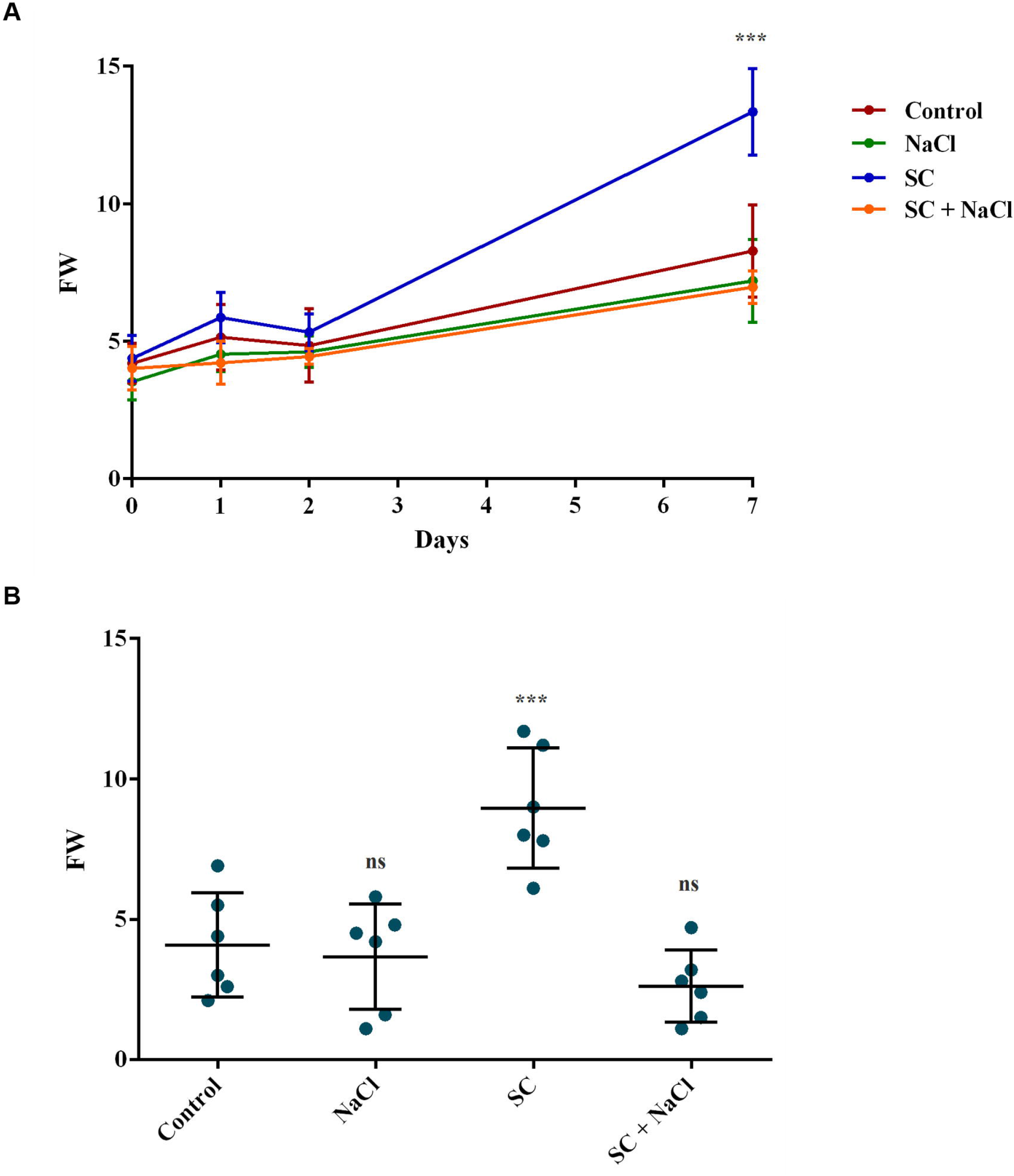
Biomass quantification of tomato plants. (A) Fresh weight biomass accumulation of tomato plants at time 0, 1, 2, and 7 days (n= 6; mean± SD) treated with sildenafil citrate in presence (SC NaCl) and in absence (SC) of 100 rnM NaCl salt-stress, under salinity (100 mM NaCl) or in control. (B) Scatterplot of tomato fresh weight at time 7 days (n= 6; mean± SD) with the same treatment of graph A. One-way ANOVA was performed with confidential level of 95%; ***p < 0.001 vs control; Dunnett’s post hoc test).

On the other hand, the analysis of root architecture pointed out that most of the tomato roots presented a diameter lower than 1 cm (Table 1), regardless the treatment, and the range between 0.1 - 0.3 cm was the most frequent in all the cases. Surprisingly, the treatment with SC in non-stressed plants provoked an increase of 56% of the total root length compared to the control. The distribution of root length over diameter classes evide1nced that all diameter classes increased following SC treatment, even though the higher diameters showed the highest increase in length (2.2- and 1.8-folds increase for 0.8-1 and > 1 cm diameters). In contrast, root length in stressed plants did not show differences compared to the control, regardless they were treated with SC or not.

**Table 1.**
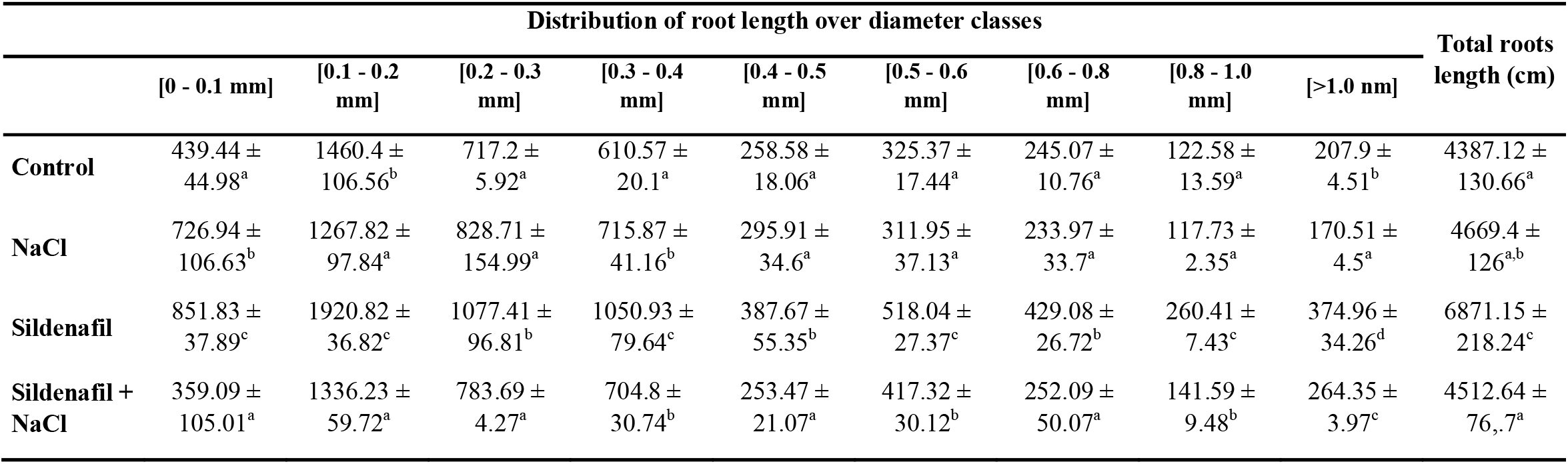
Root length distribution over different diameter classes: small (ø□=□0.0-0.2□mm), medium (ø□=□0.2–1.0□mm) and large (ø > 1.0□mm) roots diameters in each tomato plants. The values are expressed as mean values□±□standard deviation of root length (cm). Superscript letters within each column indicate differences among treatments, resulted from ANOVA (p□<□.05), Duncan’s post-hoc test.

|  | Distribution of root length over diameter classes |  |  |  |  |  |  |  |  | Total roots length (cm) |
| --- | --- | --- | --- | --- | --- | --- | --- | --- | --- | --- |
|  | [0 - 0.1 mm] | [0.1 - 0.2 mm] | [0.2 - 0.3 mm] | [0.3 - 0.4 mm] | [0.4 - 0.5 mm] | [0.5 - 0.6 mm] | [0.6 - 0.8 mm] | [0.8 - 1.0 mm] | [>1.0 mm] |  |
| **Control** | 439.44 ± 44.98<sup>a</sup> | 1460.4 ± 106.56<sup>b</sup> | 717.2 ± 5.92<sup>a</sup> | 610.57 ± 20.1<sup>a</sup> | 258.58 ± 18.06<sup>a</sup> | 325.37 ± 17.44<sup>a</sup> | 245.07 ± 10.76<sup>a</sup> | 122.58 ± 13.59<sup>a</sup> | 207.9 ± 4.51<sup>b</sup> | 4387.12 ± 130.66<sup>a</sup> |
| **NaCl** | 726.94 ± 106.63<sup>b</sup> | 1267.82 ± 97.84<sup>a</sup> | 828.71 ± 154.99<sup>a</sup> | 715.87 ± 41.16<sup>b</sup> | 295.91 ± 34.6<sup>a</sup> | 311.95 ± 37.13<sup>a</sup> | 233.97 ± 33.7<sup>a</sup> | 117.73 ± 2.35<sup>a</sup> | 170.51 ± 4.5<sup>a</sup> | 4669.4 ± 126<sup>a,b</sup> |
| **Sildenafil** | 851.83 ± 37.89<sup>c</sup> | 1920.82 ± 36.82<sup>c</sup> | 1077.41 ± 96.81<sup>b</sup> | 1050.93 ± 79.64<sup>c</sup> | 387.67 ± 55.35<sup>b</sup> | 518.04 ± 27.37<sup>c</sup> | 429.08 ± 26.72<sup>b</sup> | 260.41 ± 7.43<sup>c</sup> | 374.96 ± 34.26<sup>d</sup> | 6871.15 ± 218.24<sup>c</sup> |
| **Sildenafil + NaCl** | 359.09 ± 105.01<sup>a</sup> | 1336.23 ± 59.72<sup>a</sup> | 783.69 ± 4.27<sup>a</sup> | 704.8 ± 30.74<sup>b</sup> | 253.47 ± 21.07<sup>a</sup> | 417.32 ± 30.12<sup>b</sup> | 252.09 ± 50.07<sup>a</sup> | 141.59 ± 9.48<sup>b</sup> | 264.35 ± 3.97<sup>c</sup> | 4512.64 ± 76.7<sup>a</sup> |

### 3.2 Effect of the phosphodiesterase inhibitor on plant metabolomic profile

In this study, an untargeted metabolomics approach was used to unravel the metabolite changes triggered by cyclic nucleotide monophosphates in tomato. Overall, more than 3400 compounds were putatively annotated using the comprehensive database PlantCyc 12.6. The entire list of compounds annotated, their composite mass spectra (mass and abundance combinations), together with the compounds confirmed by MS/MS, are listed as supplementary material (Table S1). The application of the inhibitor of phosphodiesterases induced a broad reprogramming of metabolite profiles and impacted several biosynthetic pathways. At the same time, the metabolic profile was obviously affected by the NaCl treatment, and distinct responses to the stress were observed when SC was applied.

The effect of treatments was independently processed within each time-point. The unsupervised HCA was first performed to identify similarities/dissimilarities among the treatments based on their metabolic signatures. This analysis suggested that the metabolic profile of tomato was significantly influenced by SC and, to a lesser extent, by NaCl treatment (Figure 2). The stronger differences could be evidenced at the last sampling point (7 days). Therefore, this time point was further investigated to unravel the metabolic consequences of phosphodiesterase inhibition and their implication(s) in the response to salt stress. Orthogonal projection to latent structures discriminant analysis (OPLS-DA) clearly separated the different treatments from the control (Figure 3). The OPLS-DA confirmed that, despite the prevailing effect of SC, NaCl also modified the metabolic profiling, in agreement with HCA. The model was validated by the goodness-of-fit (R2Y > 0.99), the prediction ability (Q2Y > 0.9), and by the cross-validated analysis of variance (CV-ANOVA, p<0.01). The following VIP analysis was applied to point out the discriminant compounds, resulting in 213 metabolites (VIP score > 1.20). Among them, phenylpropanoids, including flavonoids and their precursors, seemed to play an important role in separation. Moreover, terpenes and the terpene hormone gibberellins were also well presented. Among hormones, jasmonic acid derivatives and auxins were also affected by the treatments. Nitrogen-containing metabolites (amino acids, amines, indoles and alkaloids) were also found as discriminant in the OPLS-DA model together with lipids and fatty acids. Interestingly, glutathione-mediate detoxification compounds were also presented as VIP markers. The list of VIP markers is provided as Supplementary Table S2.

**Figure 2.**
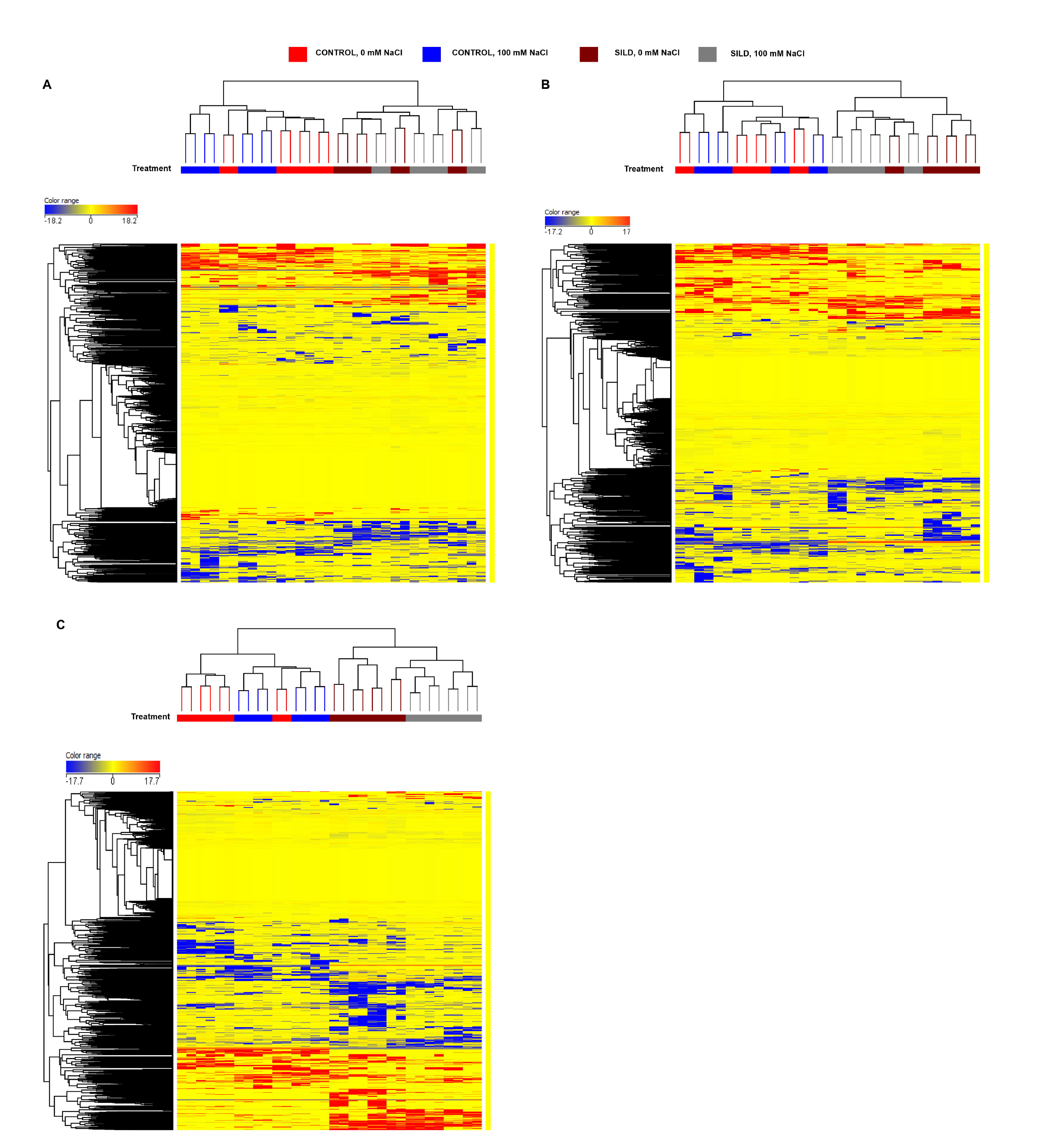
Unsupervised hierarchical cluster analysis (Euclidean distance; linkage rule: Ward) carried out from tomato leaves chemical profiles treated with SC alone or in combination with NaCl at 1 day (A), 2 days (B) and 7 days (C). Metabolites were obtained by UHPLC-ESI/QTOF-MS untargeted analysis, and their intensities used to build up the fold-change heatmap here provided.

**Figure 3.**
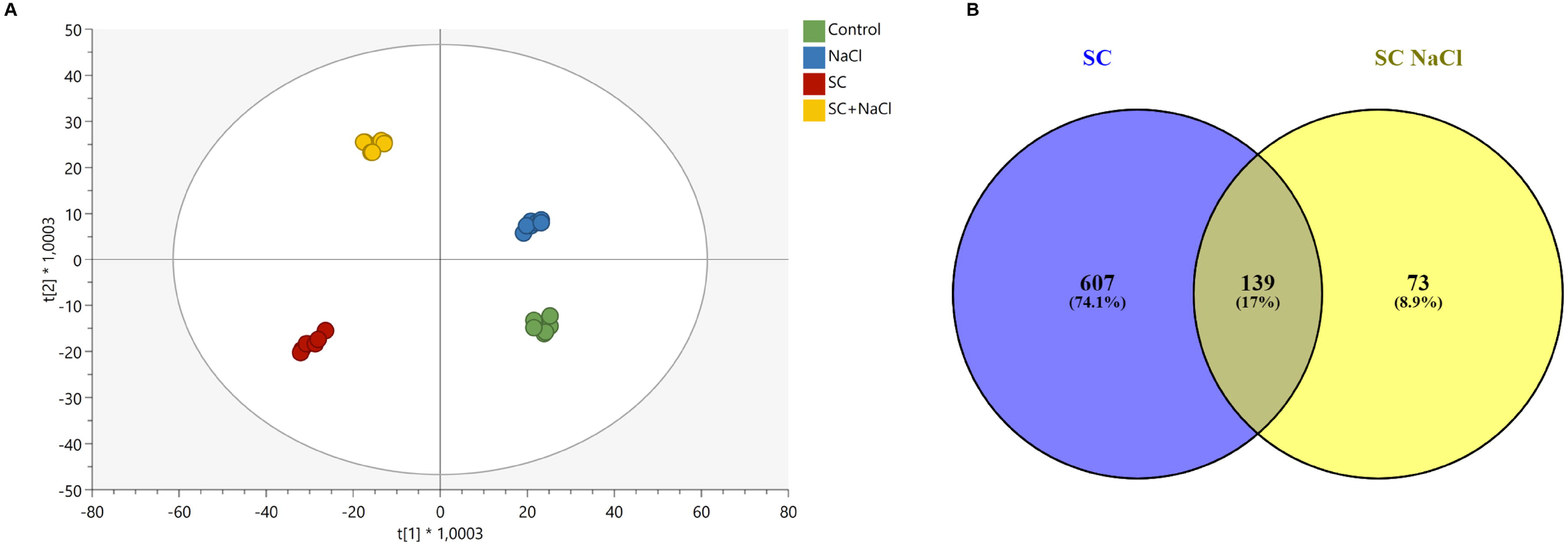
Score plot of Orthogonal projection to latent structures discriminant analysis (OPLS-DA) supervised modelling carried out from untargeted metabolomics profiles of tomato leaves treated with SC alone or in combination with NaCl at 7 days (R2Y = 0.99, Q2Y = 0.9) (A). **Supplementary Table 5**. Venn diagram comparing the differential metabolites resulting from the Volcano analysis (p<0.05; FC ≥ 2) from tomato leaves of plants treated with SC either under non-stress and 100 mM NaCl salinity conditions.(B).

#### 3.2.1. The effect of SC on nucleotides monophosphate

The application of the phosphodiesterase inhibitor resulted in a significant and generalized modulation of cyclic nucleotides monophosphate, with cGMP and cUMP showing a down-accumulation fold-change (FC) of -4.28 and -10,64 respectively. On the contrary, an up accumulation of cCMP (FC +20.60) and non-cyclic GMP (FC +9.11) could be observed. Among the other non-cyclic nucleotide monophosphate, AMP and dAMP were down accumulated (FC - 8.39 and -16.68, respectively). When salt stress conditions are considered, only cUMP could be identified by Volcano Plot analysis, showing a slight up-accumulation trend irrespectively from the application of the phosphodiesterase inhibitor (+0.79 and +0.76 in salinity and salinity + SC, respectively).

#### 3.2.2. The effect of SC on biochemical signature of tomato leaves

The effect of SC was investigated in non-stressed tomato plants by identifying the significant compounds modulated in response to phosphodiesterase inhibition. In agreement with multivariate statistics, SC largely modified the metabolomic profile of tomato plants, since over 750 compounds were significantly altered by the treatment, compared to the control (Supplementary Table S3). Figure 4 summarizes the differential metabolites, classified according to their role in the biosynthetic processes, as provided by the Omic Viewer of PlantCyc.

**Figure 4.**
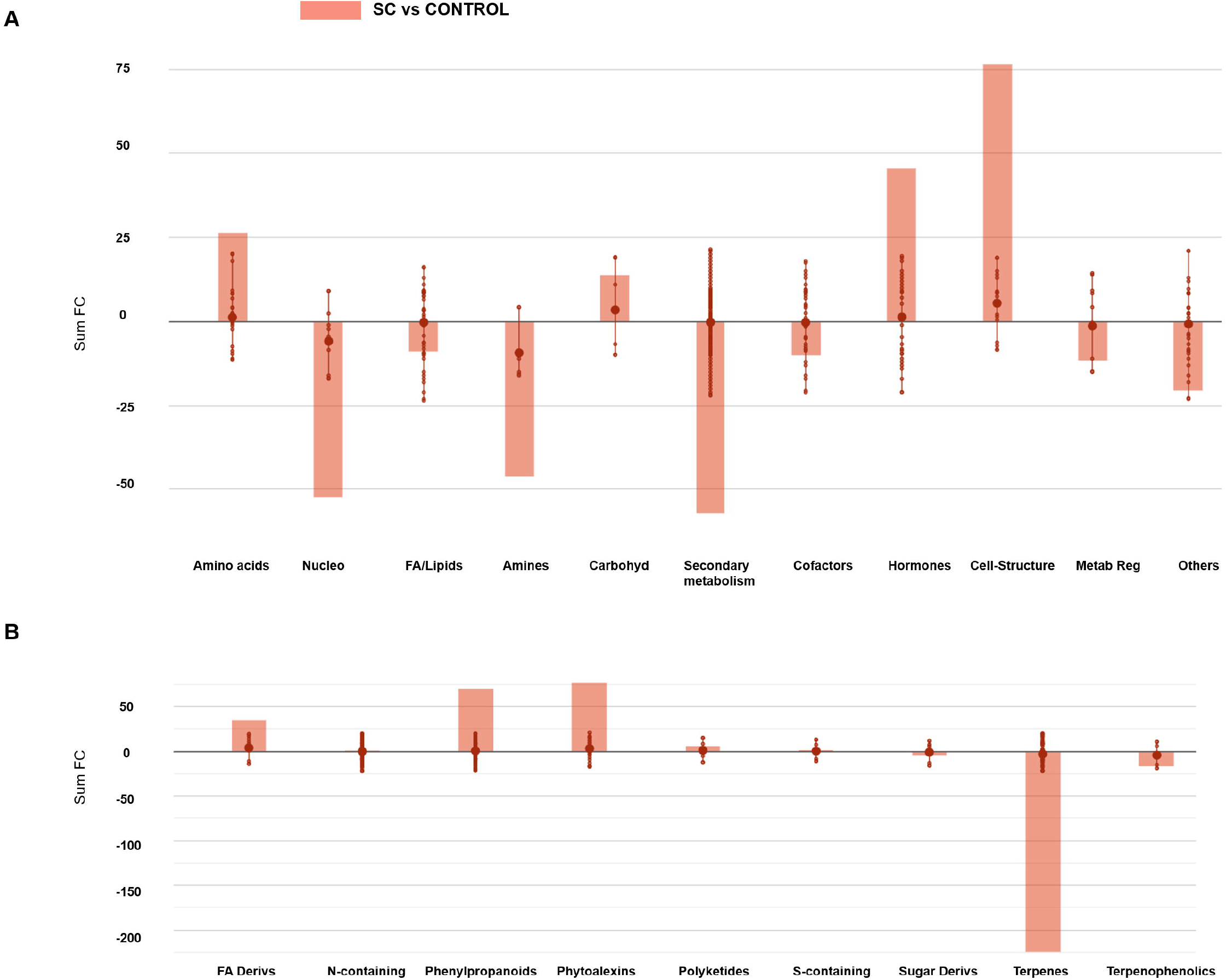
Metabolic processes (A) and secondary metabolism (B) impaired by the SC treatment. Differential metabolites and their fold-change values were elaborated using the Omic Viewer Dashboard of the PlantCyc pathway Tool software (www.pmn.plantcyc.com). The large dots represent the average (mean) of all log Fold-change (FC) for metabolites, and the small dots represent the individual log FC for each metabolite. The x-axis represents each set of subcategories, while the y-axis corresponds to the cumulative log FC. Nucleo: nucleosides and nucleotides; FA/Lipids: fatty acids and lipids; Amines: amines and polyamines; Carbohyd: carbohydrates; Cofactors: cofactors, prosthetic groups, electron carriers, and vitamins; FA Derivs: fatty acid derivatives; Nitrogen-containing: N-containing secondary metabolites; S-containing: Sulfur-containing secondary metabolites; Sugar Derivs: sugar derivatives.

Overall, SC-treated plants showed a suppression of secondary metabolism, as a generalized response. This response mainly included terpene metabolites, which exhibited a strong down-accumulation. However, an opposite trend was observed for alkaloids and phenylpropanoids that were mostly up accumulated. In particular, flavonoids and their conjugates were the most elicited among phenylpropanoids, although several cyanidin conjugates were down accumulated. Together with alkaloids, also glucosinolate-related compounds were affected by the SC treatment. In particular, the glucosinolates derived from homo- and pentahomomethionine were mostly repressed, while those derived from dihomomethionine and tryptophan were stimulated. Interestingly, the interpretation of differential metabolites highlighted also the detoxification process mediated by glutathione, the glutathione conjugates, among the processes largely affected by SC. Furthermore, other metabolites related to oxidative stress such as γ-L-glutamyl-L-cysteine and oxo-GMP (both accumulated following SC treatment), and ascorbate (being down accumulated) were found among significant compounds. Interestingly, several metabolites related to nitric oxide biosynthesis, namely L-citrulline and N-ω-hydroxy-L-arginine (both precursors of NO, both up-accumulated in SC-treated plants), the cofactor tetrahydrobiopterin, glycine betaine (up accumulated) and its precursors, as well as the NO inducer tricaffeoyl spermidine were found among differential metabolites. Similarly, the precursors of NO-mediated modification of plant cell structure occurring in hypersensitive response (*i*.*e*., the monolignols coniferaldehyde and synapaldehyde) were strongly stimulated in SC-treated plants. The concurrent modulation of salicylate 2-*O*-β-*D*-glucoside (a more soluble form of salicylate, typically involved in vacuole storage) and salicyl alcohol, both strongly up-accumulated, together with the down accumulation of salicylaldehyde and methylsalicylate can be also related to NO-mediated hypersensitive response.

It is worth mentioning that also hormones presented a significant imbalance following SC application, including several auxin precursors and auxin conjugates that were accumulated following application of SC. A similar trend was observed for cytokinins (kinetin, isopentenyladenine-9-*N*-glucoside, trans-zeatin and benzyladenine-7-*N*-glucoside, among others), while the terpene-derived hormones brassinosteroids were generally down accumulated. Concerning gibberellins, methyl GA9 and the bioactive GA7 were increased, whereas other gibberellins were down accumulated. Abscisic acid was also involved in the response to SC, with its degradation products (phaseic acid and 7-hydroxy-abscisate) being accumulated and precursors showing an opposite trend.

On the other hand, metabolites related to photosynthesis were impaired by SC application. A modulation of both chlorophyll biosynthesis and degradation intermediates was observed following treatment with SC. Despite the accumulation of both chlorophyll a and b, the flux of carbon in the chlorophyll biosynthetic pathway indicated the accumulation of downstream metabolites, in particular the degradation products of chlorophyll a. Compounds such as primary fluorescent chlorophyll catabolite, red chlorophyll catabolite or pheophorbides increased at the expense of the compounds upstream.

#### 3.2.3. The effect of SC on biochemical signatures of tomato leaves under salt stress conditions

The significant metabolites impaired by the SC in tomato under salinity (provided in Table S4) were interpreted by means of the Pathway tool analysis as previously reported for non-stressed plants. More than 200 compounds were modulated by either NaCl or SC treatment under salinity, with secondary metabolites being particularly affected. Figure 5 summarizes the differential metabolites involved SC-treated tomato plants in response to salinity, classified according to their role in the biosynthetic processes as provided by the Omic Viewer of PlantCyc. Overall, salinity elicited secondary metabolism, while a decrease was observed when SC was applied under salinity. Phenylpropanoids and terpenes increased and decreased, respectively, in the presence of both NaCl and SC-NaCl, compared to the control. In more detail, flavonoid conjugates were accumulated in NaCl-treated plants, while an unclear trend was observed for SC-NaCl. Although several flavonoids conjugates also increased in the presence of SC-NaCl, compounds such as 3-(4-hydroxyphenyl)pyruvate, pterostilbene or 4-coumaryl alcohol were stimulated in these treated plants. Concerning N-containing secondary metabolites, alkaloids and glucosinolate related compounds accumulated under NaCl whereas a decrease was observed for SC-NaCl.

**Figure 5.**
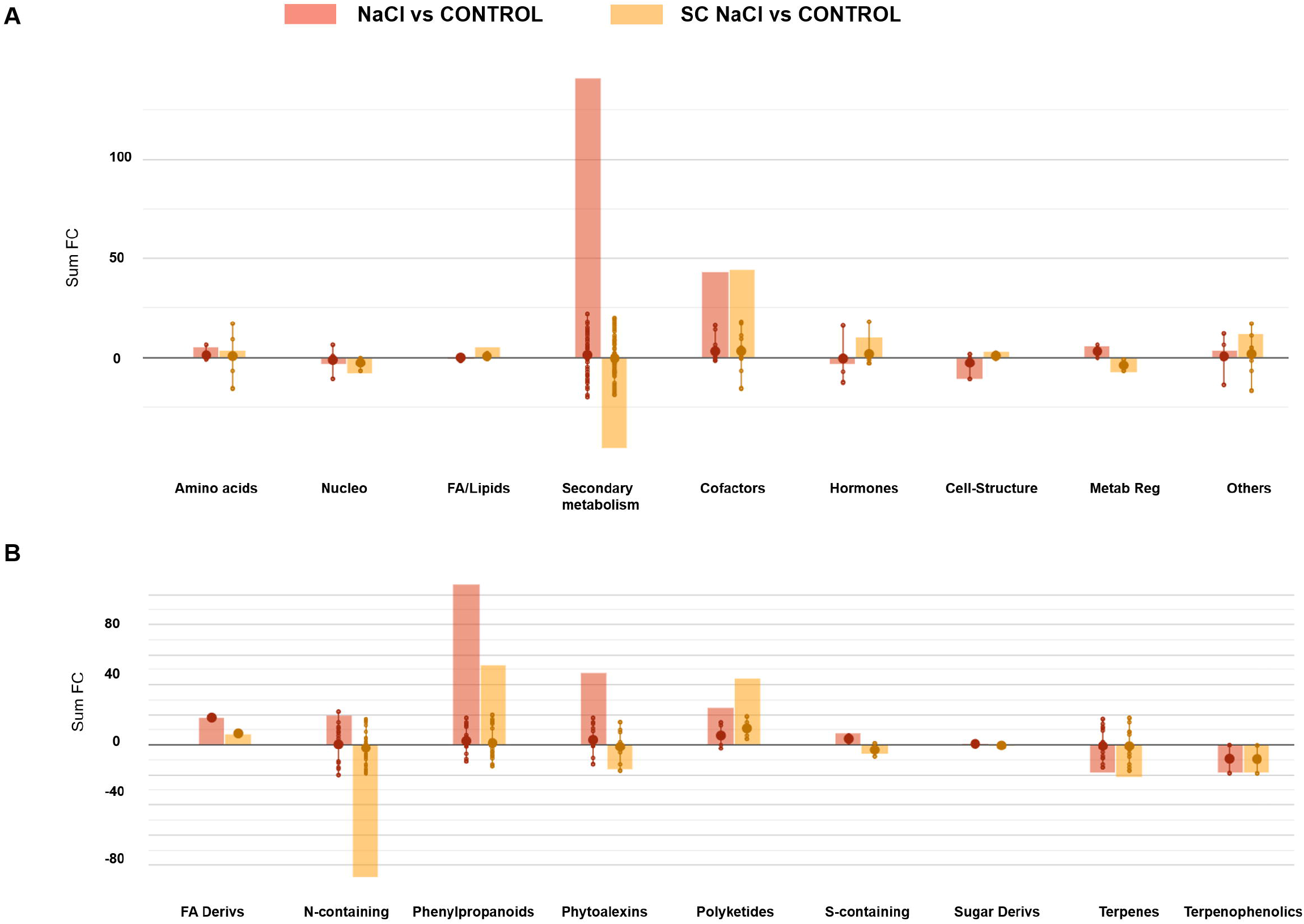
Metabolic processes (A) and secondary metabolism (B) impaired by the NaCl stress alone or in combination with SC. Differential metabolites and their fold-change values were elaborated using the Omic Viewer Dashboard of the PlantCyc Pathway Tool software (www.pmn.plantcyc.com). The large dots represent the average (mean) of all log Fold-change (FC) for metabolites, and the small dots represent the individual log FC for each metabolite. The x-axis represents each set of subcategories, while the y-axis corresponds to the cumulative log FC. Nucleo: nucleosides and nucleotides; FA/Lipids: fatty acids and lipids; Amines: amines and polyamines; Carbohyd: carbohydrates; Cofactors: cofactors, prosthetic groups, electron carriers, and vitamins; FA Derivs: fatty acid derivatives; Nitrogen-containing: N-containing secondary metabolites; S-containing: Sulfur-containing secondary metabolites; Sugar Derivs: sugar derivatives.

The accumulation of metabolites involved in chlorophyll biosynthesis was induced by SC application under salinity stress, while an opposite trend was observed for NaCl alone. In this sense, several porphyrin and organometallic compound (*i*.*e*., epoxypheophorbide, pheophorbide, protoporphyrinogen and oxo-magnesium-protoporphyrin-monomethyl ester) increased in SC-NaCl and decreased in NaCl treatment suggesting a specific mechanism triggered by SC, irrespective of the stress conditions. Regarding hormones, similar patterns were found in both SC and SC-NaCl-treated plants. Gibberellins and their precursors were down-accumulated in NaCl and, in a lesser extent, in SC-NaCl. Abscisic aldehyde was strongly down-accumulated in SC-NaCl and ascorbate, auxin conjugates and cytokinins were up-accumulated in both treatments.

A venn analysis between SC and SC-NaCl was then carried out to compare treatments, confirming a hierarchically more pronounced effect of SC than NaCl (Supplementary Table S5), in agreement with multivariate statistics. Among the 212 compounds impaired by the NaCl stress, 139 were in common with the SC treatment. The decrease of terpene compounds overlapped for both treatments.

## 4. DISCUSSION

There is a fast-growing interest about the role played by cyclic nucleotide monophosphates in plant growth and response to environment. At the sight of the little information about cNMP in plants and taking into account the evidences of their crucial role in plant signalling and development, we tested an inhibitor of PDE to highlight the role of these molecules in the biochemical fingerprints of tomato plants.

It is consolidated that SC inhibits the PDE5 in humans, which specifically catalyses the hydrolysis of cGMP to GMP (Turko *et al*., 1999). However, in plants, two principal classes of PDE have been described: those that catalyse the hydrolysis of 2′,3′-cNMP to 2′-NMP and those involved in the hydrolysis of 3′,5′-cNMPs to produce a mixture of 3′-NMP and 5′-NMP. Enzymes with a dual enzymatic function have been identified in *Solanaceae* and *Fabaceae* (Swiezawska *et al*., 2018). (Kasahara *et al*., 2016) reported the MpCAPE gene, derived from AC gene in *Marchantia polymorpha* with both C-terminal AC catalytic domain similar to those of class III ACs and an N-terminal cyclic nucleotide PDE. The analysis of recombinant proteins showed PDE activities hydrolysing both cAMP and cGMP. However, cAMP was a much more favourable substrate than cGMP (Kasahara *et al*., 2016). Abel *et al*., (2000) also observed that extracellular phosphodiesterases derived from tomato cell cultures greater hydrolysed 2′-3′-cyclic NMP to 3′-NMP than 3′-5′-cyclic isomers to a 3′-NMP and 5′-NMP, regardless the nucleobase. Both cNMP and their inactive forms were well represented as markers of the SC treatment. Even if PDE are not fully characterized in plant, and the specific effect of SC on different PDE is unknown, distinct and significantly different profiles could be observed regarding NMP in tomato leaves. Although information about the half-life of cNMP in plants is unknown, we can assume they are short living compounds in the cell, compared to other eukaryotic cells like animals. This may explain the complex and distinctive trends we observed for the different cNMP. Nonetheless, it is clear that SC-mediated PDE inhibition was more evident for cCMP, compared to cGMP and cUMP. This may indicate that SC provided different degrees if inhibition towards the different PDE in tomato.

Still, SC application impacted the accumulation of several NMP and, due to their crucial implication in biological processes, this resulted in the profound shaping of metabolomic profiles we could observe. In fact, irrespective from the specific degree of inhibition of SC towards the different PDEs, multivariate statistics clearly highlighted that the treatment induced an evident modulation of tomato metabolomic profiles. Given their function as secondary messengers, there are clear evidences of cNMP role in key physiological processes such as the cation fluxes, photosynthesis and stomatal opening (Gehring and Turek, 2017). Therefore, it is hardly surprising that the disruption of cNMP profile provokes critical changes in metabolic profiles. In fact, we observed a modulation of chlorophyll pigments following the addition of SC in both non-stressed and stressed plants. Thomas et al. (2013) suggested that cAMP might be involved in the photosynthesis since several differentially expressed proteins with a role in the photosystem II were impaired by the addition of cAMP (Thomas *et al*., 2013). Segovia *et al*., (2001) observed a direct relationship between cAMP accumulation and photosynthesis in the red macroalga *Porphyra leucosticte* (Segovia *et al*., 2001). In more detail, an interplay between cAMP and photosynthesis has been proposed, where photosynthetic electron transport was involved in the regulation of cAMP accumulation in *Dictyota dichotoma* and *Gelidium sesquipedale* (Gordillo *et al*., 2004).

The signal cascade of cNMP is correlated with the activity of downstream effectors, i.e. protein kinases (PKN), ion channels, transcription factors (Swiezawska *et al*., 2018), as well as cyclic Nucleotide-Binding Proteins (CNBPs). CNBPs described in *A. thaliana* might have a role in photorespiration and Calvin cycle (Donaldson *et al*., 2016). These authors proposed a crosstalk between cNPM, NO and H_2_O_2_ signalling for the modulation of these proteins. For instance, NO stimulates the guanylyl cyclase (GC) to produce cGMP, and cAMP has been proposed to modulated oxidative stress by the reduction of Ca^2+^ influx and K^+^ efflux via CNBPs (Blanco *et al*., 2020). Several metabolomic clues indicate that the NO signalling cascade was significantly altered by the treatment with SC. In more detail, the increase of phenylpropanoids is known to be elicitated by NO through the modulation of Phenylalanine Ammonia Lyase (PAL) and cinnamic acid-4-hydroxylase (C4H) (Romero-Puertas *et al*., 2004). Nonetheless, the inhibition of jasmonate accumulation (Zhou *et al*., 2015), the alteration of polyamines (Tun *et al*., 2006), biopterin-related compounds, salicylate and glutathione conjugates (Romero-Puertas *et al*., 2004), as well as the elicitation and glycine betaine (Ullah *et al*., 2016) *i*.*e*. a significant part of the differential responses we could observe through metabolomics, can be all related to the NO-mediated signalling cascade. Furthermore, it is known that NO has a pivotal role in triggering plant hypersensitive response, a process involving salicylate and ascorbate to finally modulate ROS. The increase of monolignols we observed, can be related to the increased cell wall lignification occurring during hypersensitive response (Romero-Puertas *et al*., 2004). An appropriate balance between ROS and NO production is actually required in hypersensitive response (Delledonne *et al*., 2001). Moreover, NO governs ROS balance from one side by eliciting the production of superoxide anions O_2_^-^ via NADPH Oxidase, and from the other side by providing a negative feedback limiting ROS by S-nitrosylation of NADPH Oxidase (Yun *et al*., 2011). As a consequence, NO and ROS signalling are tightly interconnected and are both part of a pivotal crosstalk in plant. Together with ascorbate and salicylate, several other molecules related to ROS scavenging and detoxification processes were shaped by SC treatment. For instance, folates, which increased in the presence of SC, are linked to ROS detoxification (Gorelova *et al*., 2017). Similarly, 8-oxo-GMP (a marker of oxidative imbalance) and glutathione conjugates were impaired by SC, strengthening the involvement of cNMP in the NO and ROS crosstalk. Consistently, Paradiso *et al*., (2020) recently determined that the minimization of intracellular cAMP by means of a cAMP-sponge caused the accumulation of reactive oxygen species and impaired ROS-related enzymes such as ascorbate peroxidase. On the other hand, Nitric Oxide Synthase□3 is post-translationally regulated by a cGMP dependent protein kinase in endothelial cells of animals (John *et al*., 2006). Furthermore, human Nitric Oxide Synthases seems to be regulated by calmodulin binding following an increase of cytosolic Ca^2+^ concentration, a process involving CNGC (Piech *et al*., 2003). The balance between Ca^2+^ ions and cNMP seems to be crucial for cyclic nucleotide gated channels, cyclase activity and for specific signaling pathways (Swiezawska-Boniecka *et al*., 2021). On the other hand, it is now recognized that distinct signal transduction systems have evolved to allow adaptive responses in plant, and cNMP have been related to this signaling complementation (Swiezawska *et al*., 2018). A summary of the cNMP, NO and ROS signal interplay we postulated from our metabolomics insights is provided in figure 6. Despite future confirmation in plants are to be provided, a similar cross-regulation can be proposed in our experiments.

**Figure 6.**
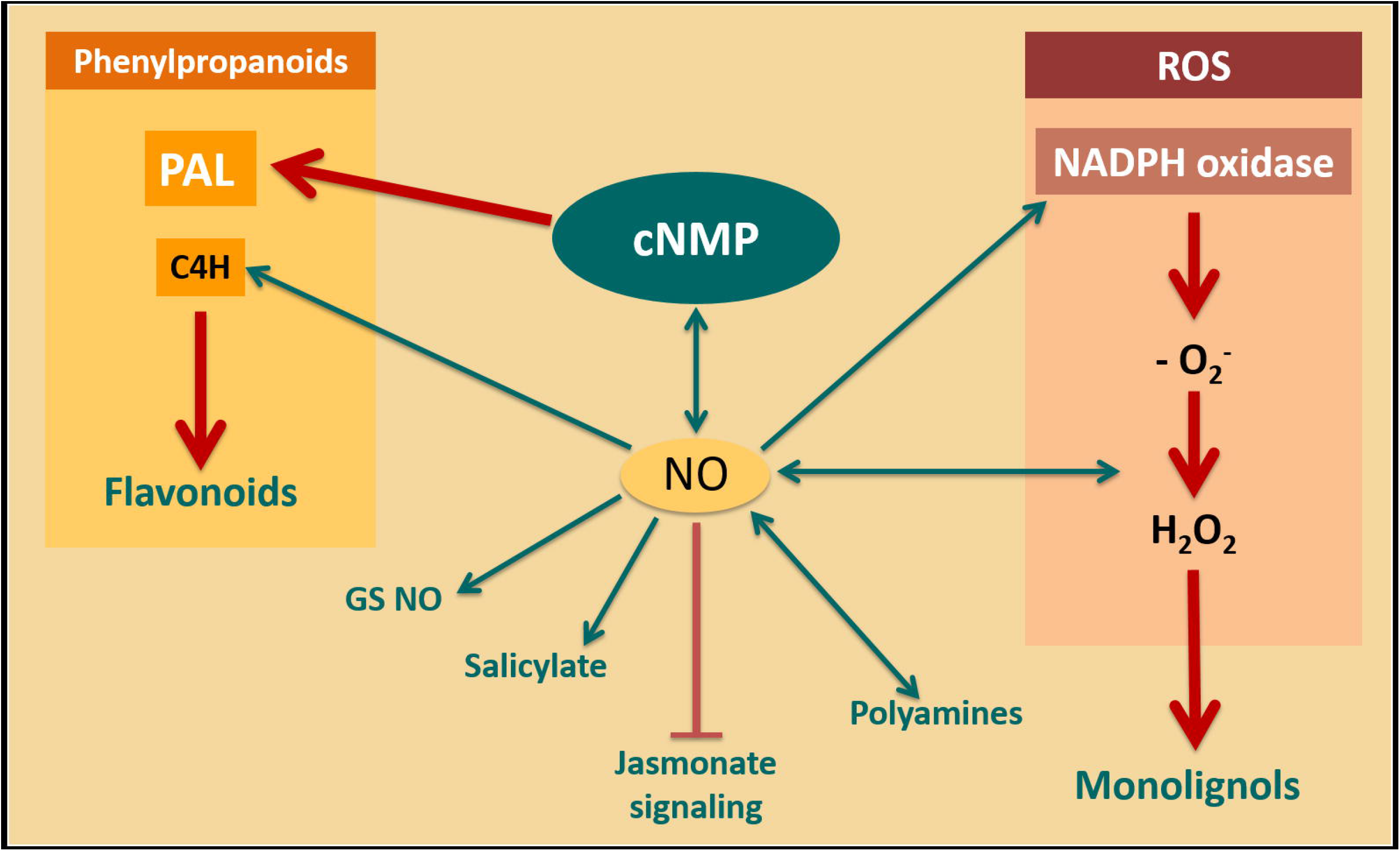
A summary of the integration of cNMP, NO and ROS signaling as postulated in tomto from metabolomics of plants treated with a phosphodiesterase inhibitor.

Together with the modulation of cellular signalling processes, an interactions between cNMP and phytohormones has been also observed (Gehring and Turek, 2017). Our findings confirmed the involvement of several hormones related to plant growth and development. Auxins were strongly impaired in tomato leaves by the addition of SC, with several precursors being accumulated and inactivation forms being decreased. Auxins are the main hormones regulating root development through the interplay of local biosynthesis, transport, perception and signalling (Olatunji *et al*., 2017). Not only cGMP and cAMP but also other secondary messenger molecules such as NO, ROS, Ca^2+^ or diacylglycerol are involved in auxin perception and signalling. For instance, cAMP-regulated guanine nucleotide exchange factors directly relate the auxin signal transduction. Similarly, cGMP affects auxin-regulated gene expression and auxin/IAA degradation by regulating PKN activity (Di *et al*., 2015). Recently, auxins have been associated to the modulation of chlorophyll and ROS scavenging (Salazar-Iribe and De-la-Peña, 2020). The increase of auxins in SC-treated plants might also explain their higher root development, in agreement with previous findings. Indeed, it was reported that cGMP and NO are messengers involved in the auxin-induced increase of adventitious roots (Pagnussat *et al*., 2003). The concurrent increase in cytokinins following SC treatment, the primary interaction partners of auxins, also supports the increase in biomass we observed. Among the several roles cytokinins have in plant, the promotion of cell division is likely the best known. The balance between auxins and cytokinins has a pivotal importance in the control of shoot and root apical meristems (Su *et al*., 2011) via a cross-regulated loop mechanism (Kotov and Kotova, 2018). Besides, an increase of the active gibberellin GA7 was observed in our treated tomato plants. Still referring to the increase in biomass we observed, it can be noted that gibberellins are involved in cell elongation processes, and cGMP-dependent protein kinases are a fundamental component of gibberellins signalling (Shen *et al*., 2019).

Concerning the response of tomato plant to SC application under salinity conditions, an effect at metabolite level could be observed even though biomass was not affected. In this regard, several studies associated the c-AMP signalling with osmotic stress, through the modulation of Na^+^ flux or phosphorylation of a plant aquaporins by a cAMP-dependent protein kinases (Maathuis and Sanders, 2001; Vera-Estrella *et al*., 2004).

From a wider perspective, cNMP are also involved in plant response to adverse conditions (Blanco *et al*., 2020). The phenylpropanoids pathway was among the processes undergoing the more evident impairment following the addition of SC. Chlorophyll intermediates were decreased under salinity, a key process known to be displaced by plants in order to reduce the accumulation of ROS from the photosynthetic electron transport chain (Foyer and Noctor, 2005). Interestingly, when SC was applied, an opposite trend could be observed. This may indicate a better ability of tomato to cope with salinity-related oxidative imbalance and acclimation to abiotic stress (Foyer and Noctor, 2009). Looking at hormone profiles, similar trends could be observed for auxins and cytokinins between salinity or non-saline conditions, whereas gibberellins showed opposite trends. Despite the NO cascade has been reviewed as a promoter of plant growth under salinity (Nabi *et al*., 2019) the exogenous PDA inhibitor did not result in an increase of tomato biomass in our experiments. These discrepancies could be partially ascribed to the different responses triggered by NO, that involves a broad cascade not limited to cNMP, and/or to the fact that NO and ROS interplay was reported to relieve the early response to salt stress in soybean (Dinler *et al*., 2014) whereas a later response was investigated in this study. Nonetheless, different metabolic responses related to oxidative stress (ascorbate, phenylpropanoids and chlorophyll intermediates level) were modulated by SC in tomato under salinity. Similarly, endogenous NO, ascorbate and salinity tolerance have been related in pepper (Kaya *et al*., 2020). Starting from these findings, more targeted studies are suggested to better unravel the role of cNMP in plant tolerance to salinity, likely including early and late responses, the whole NO cascade, enzymes activity (*e*.*g*., the Foyer-Halliwell-Asada cycle) and different omic approaches. Indeed, the interesting premises relating to the cNMP role in relieving oxidative stress may be very important to identify cultivars or treatments able to support plant growth in saline soils.

## 5. CONCLUSIONS

The understanding about the role played by cNMP in plants is a novel topic that finds scientific and practical interest. In this study, we used SC, an exogenous PDE inhibitor, to highlight the signalling cascade mediated by cNMP. The effect of inhibiting PDE seemed to affect not only cGMP, as expected, but also others cNMP including cAMP. The alteration of cNMP provoked a broad metabolic reprogramming, leading to an impairment of several pathways to include both primary metabolism processes (as photosynthesis) and secondary metabolism. Biomass significantly and consistently increased following the application of SC, and a modulation of root architecture was observed. Metabolomics pointed out a complex crosstalk network of phytohormones and other signalling compounds, together with redox imbalance processes, with NO being suggested as a main driver of plant response to treatments. Salinity conditions damped down the growth-promoting effects of SC, but indicated interesting implications in plant mitigation to stress-related detrimental effects.

## Abbreviations

ANOVA: analysis of variance
cAMP: cyclic AMP (adenosine 3′:5′-cyclic monophosphate)
cCMP: cyclic CMP (cytidine, 3′:5′-cyclic monophosphate)
cdTMP: cyclic dTMP (2′-deoxythymidine 3′:5′-cyclic monophosphate)
cGMP: cyclic GMP (Guanosine 3′:5′-cyclic monophosphate)
cIMP: cyclic IMP (inosine 3′:5′-cyclic monophosphate)
CNGC: cyclic nucleotide gated channel
cNMP: cyclic nucleotide monophosphate
COSMOS: COrdination of Standanrds in MetabOlomicS
CV-ANOVA: cross-validated analysis of variance
cUMP: cyclic UMP (uridine 3′:5′-cyclic monophosphate)
dAMP: deoxyadenosine monophosphate
DW: dry weight
FC: fold-change
FW: fresh weight
FWHM: full width half maximum
HCA: hierarchical cluster analysis
IAA: indoleacetic acid
MS: mass spectrometry
NADPH: nicotinamide-adenine dinucleotide phosphate
NO: nitric oxide
OPLS-DA: orthogonal projection to latent structures discriminant analysis
PAL: phenylalanine ammonial lyase
PDE: Phosphodiesterase
PKA: protein kinase A
PKG: protein kinase G
QC: quality control
ROS: reactive oxygen species
SC: sildenafil citrate
UHPLC/QTOF-MS: ultra high-performance liquid chromatography -quadrupole time-of-flight mass spectrometry
VIP: variables importance in projection

## ACKNOWLEDGEMENTS

This work is the result of a postdoctoral contract for the training and improvement abroad of research staff (Begoña Miras-Moreno; 21252/PD/19) financed by the Consejería de Empleo, Universidades, Empresa y Medio Ambiente of the CARM, through the Fundación Séneca-Agencia de Ciencia y Tecnología de la Región de Murcia (Spain). The authors are grateful to prof Giuseppe Colla, University of Tuscia for his kind support in root morphology analysis. The authors wish to thanks the “Romeo ed Enrica Invernizzi” foundation, Milan (Italy) for its kind support to the metabolomics facility.

## AUTHOR CONTRIBUTION

Conceptualization: L.L., B.M.M.; Data Curation: L.Z., B.M.M.; Formal Analysis: B.M.M., L.Z., B.S.; Funding Acquisition: L.L.; Investigation: B.M.M., L.Z., B.S.; Methodology: L.L., B.M.M.; Supervision: L.L.; Writing – Original Draft Preparation: L.L., B.M.M., L.Z., B.S.; Writing – Review & Editing: L.L.

## SUPPLEMENTARY MATERIAL

**Supplementary Table 1**. Whole dataset produced from untargeted metabolomics carried out in tomato leaves treated with SC either under stressed (100 mM NaCl) and non-stress conditions. Compounds are presented with individual intensities and composite mass spectra.

**Supplementary Table 2**. Discriminant metabolites identified by the variable importance in projection (VIP) analysis following OPLS-DA modelling of metabolome in tomato leaves treated with SC, either under non-stressed and stress (100 mM NaCl) conditions during a 7-days trial. Compounds were selected as discriminant by possessing a VIP score>1.20.

**Supplementary Table 3**. Differential metabolites derived from Volcano analysis (*p*<0.05; FC ≥ 2) in leaves of tomato treated with SC, after 7 days.

**Supplementary Table 4**. Differential metabolites derived from Volcano analysis (*p*<0.05; FC ≥ 2) in leaves of tomato treated with SC under 100 mM NaCl salinity, after 7 days.

## REFERENCES

Abel S, Nürnberger T, Ahnert V, Krauss GJ, Glund K. 2000. Induction of an extracellular cyclic nucleotide phosphodiesterase as an accessory ribonucleolytic activity during phosphate starvation of cultured tomato cells. Plant Physiology 122, 543–552.

Arazi T, Kaplan B, Fromm H. 2000. A high-affinity calmodulin-binding site in a tobacco plasma-membrane channel protein coincides with a characteristic element of cyclic nucleotide-binding domains. Plant Molecular Biology 42, 591–601.

Blanco E, Fortunato S, Viggiano L, de Pinto MC. 2020. Cyclic amp: A polyhedral signalling molecule in plants. International Journal of Molecular Sciences 21, 4862.

Caspi R, Dreher K, Karp PD. 2013. The challenge of constructing, classifying, and representing metabolic pathways. FEMS Microbiology Letters 345, 85–93.

Delledonne M, Zeier J, Marocco A, Lamb C. 2001. Signal interactions between nitric oxide and reactive oxygen intermediates in the plant hypersensitive disease resistance response. Proceedings of the National Academy of Sciences of the United States of America 98, 13454–13459.

Di DW, Zhang C, Guo GQ. 2015. Involvement of secondary messengers and small organic molecules in auxin perception and signaling. Plant Cell Reports 34, 895–904.

Dinler BS, Antoniou C, Fotopoulos V. 2014. Interplay between GST and nitric oxide in the early response of soybean (Glycine max L.) plants to salinity stress. Journal of Plant Physiology 171, 1740–1747

Donaldson L, Meier S, Gehring C. 2016. The arabidopsis cyclic nucleotide interactome. Cell Communication and Signaling 14, 10.

Duszyn M, Swiezawska B, Szmidt-Jaworska A, Jaworski K. 2019. Cyclic nucleotide gated channels (CNGCs) in plant signalling—Current knowledge and perspectives. Journal of Plant Physiology 241, 153035.

Foyer CH, Noctor G. 2005. Oxidant and antioxidant signalling in plants: A re-evaluation of the concept of oxidative stress in a physiological context. Plant, Cell and Environment 28, 1056–1071.

Foyer CH, Noctor G. 2009. Redox regulation in photosynthetic organisms: Signaling, acclimation, and practical implications. Antioxidants and Redox Signaling 11, 861–905.

Ganugi P, Miras-Moreno B, Garcia-Perez P, Lucini L, Trevisan M. 2020. Concealed metabolic reprogramming induced by different herbicides in tomato. Plant Science 303, 110727.

Gehring C, Turek IS. 2017. Cyclic nucleotide monophosphates and their cyclases in plant signaling. Frontiers in Plant Science 8, 1704.

Gordillo FJL, Segovia M, López-Figueroa F. 2004. Cyclic AMP levels in several macroalgae and their relation to light quantity and quality. Journal of Plant Physiology 161, 211–217.

Gorelova V, Ambach L, Rébeillé F, Stove C, Van Der Straeten D. 2017. Folates in plants: Research advances and progress in crop biofortification. Frontiers in Chemistry 5, 21.

Irving HR, Cahill DM, Gehring C. 2018. Moonlighting proteins and their role in the control of signaling microenvironments, as exemplified by cGMP and phytosulfokine receptor 1 (PSKR1). Frontiers in Plant Science 9, 415.

Isner JC, Maathuis FJM. 2018. CGMP signalling in plants: From enigma to main stream. Functional Plant Biology 42(2), 93–101.

John TA, Ibe BO, Usha Raj J. 2006. Oxygen alters caveolin-1 and nitric oxide synthase-3 functions in ovine fetal and neonatal lung microvascular endothelial cells. American Journal of Physiology -Lung Cellular and Molecular Physiology 291(5), 1079–93.

Kasahara M, Suetsugu N, Urano Y, et al. 2016. An adenylyl cyclase with a phosphodiesterase domain in basal plants with a motile sperm system. Scientific Reports 6, 39232.

Kaya C, Ashraf M, Alyemeni MN, Ahmad P. 2020. The role of endogenous nitric oxide in salicylic acid-induced up-regulation of ascorbate-glutathione cycle involved in salinity tolerance of pepper (Capsicum annuum L.) plants. Plant Physiology and Biochemistry 147, 10–20.

Kimura S, Sinha N. 2008. Tomato (Solanum lycopersicum): A model fruit-bearing crop. Cold Spring Harbor Protocols 3.

Kotov AA, Kotova LM. 2018. Auxin-cytokinin interactions in the regulation of correlative inhibition in two-branched pea seedlings. Journal of Experimental Botany 69, 2967-2968.

Maathuis FJM, Sanders D. 2001. Sodium uptake in Arabidopsis roots is regulated by cyclic nucleotides. Plant Physiology 127, 1617–1625.

Marondedze C, Wong A, Thomas L, Irving H, Gehring C. 2017. Cyclic nucleotide monophosphates in plants and plant signaling. Handbook of Experimental Pharmacology, 87–103.

Martinez-Atienza J, Van Ingelgem C, Roef L, Maathuis FJM. 2007. Plant cyclic nucleotide signalling: Facts and fiction. Plant Signaling and Behavior 2, 540–543.

Miras-Moreno B, Corrado G, Zhang L, et al. 2020. The metabolic reprogramming induced by sub-optimal nutritional and light inputs in soilless cultivated green and red butterhead lettuce. International Journal of Molecular Sciences 21, 6831.

Nabi RBS, Tayade R, Hussain A, Kulkarni KP, Imran QM, Mun BG, Yun BW. 2019. Nitric oxide regulates plant responses to drought, salinity, and heavy metal stress. Environmental and Experimental Botany 161, 120–133.

Olatunji D, Geelen D, Verstraeten I. 2017. Control of endogenous auxin levels in plant root development. International Journal of Molecular Sciences 18, 2584.

Pagnussat GC, Lanteri ML, Lamattina L. 2003. Nitric oxide and cyclic GMP are messengers in the indole acetic acid-induced adventitious rooting process. Plant Physiology 132, 1241–1248.

Paradiso A, Domingo G, Blanco E, Buscaglia A, Fortunato S, Marsoni M, Scarcia P, Caretto S, Vannini C, de Pinto MC. 2020. Cyclic AMP mediates heat stress response by the control of redox homeostasis and ubiquitin-proteasome system. Plant Cell and Environment 43, 2727–2742..

Paul K, Sorrentino M, Lucini L, et al. 2019. A combined phenotypic and metabolomic approach for elucidating the biostimulant action of a plant-derived protein hydrolysate on tomato grown under limited water availability. Frontiers in Plant Science 10, 493.

Piech A, Dessy C, Havaux X, Feron O, Balligand JL. 2003. Differential regulation of nitric oxide synthases and their allosteric regulators in heart and vessels of hypertensive rats. Cardiovascular Research 57, 456–467.

Romero-Puertas MC, Perazzolli M, Zago ED, Delledonne M. 2004. Nitric oxide signalling functions in plant-pathogen interactions. Cellular Microbiology 6, 795–803.

Salazar-Iribe A, De-la-Peña C. 2020. Auxins, the hidden player in chloroplast development. Plant Cell Reports 39, 1595–1608.

Salek RM, Neumann S, Schober D, et al. 2015. COordination of Standards in MetabOlomicS (COSMOS): facilitating integrated metabolomics data access. Metabolomics 11, 1587–1597.

Salmi ML, Morris KE, Roux SJ, Porterfield DM. 2007. Nitric oxide and cGMP signaling in calcium-dependent development of cell polarity in Ceratopteris richardii. Plant Physiology 144, 94–104.

Segovia M, Gordillo FJL, Schaap P, Figueroa FL. 2001. Light regulation of cyclic-AMP levels in the red macroalga Porphyra leucosticta. Journal of Photochemistry and Photobiology B: Biology 64, 69–74.

Senizza B, Rocchetti G, Ghisoni S, Busconi M, De Los Mozos Pascual M, Fernandez JA, Lucini L, Trevisan M. 2019. Identification of phenolic markers for saffron authenticity and origin: An untargeted metabolomics approach. Food Research International 126, 108584

Shen Q, Zhan X, Yang P, Li J, Chen J, Tang B, Wang X, Hong Y. 2019. Dual activities of plant cGMP-dependent protein kinase and its roles in gibberellin signaling and salt stress[OPEN]. Plant Cell 31, 3073–3091.

Su YH, Liu YB, Zhang XS. 2011. Auxin-cytokinin interaction regulates meristem development. Molecular Plant 4, 616–625.

Swiezawska-Boniecka B, Duszyn M, Kwiatkowski M, Szmidt-Jaworska A, Jaworski K. 2021. Cross Talk Between Cyclic Nucleotides and Calcium Signaling Pathways in Plants–Achievements and Prospects. Frontiers in Plant Science 12, 643560.

Swiezawska B, Duszyn M, Jaworski K, Szmidt-Jaworska A. 2018. Downstream targets of cyclic nucleotides in plants. Frontiers in Plant Science 9, 1428.

Thomas L, Marondedze C, Ederli L, Pasqualini S, Gehring C. 2013. Proteomic signatures implicate cAMP in light and temperature responses in Arabidopsis thaliana. Journal of Proteomics 83, 47–59.

Tsugawa H, Cajka T, Kind T, Ma Y, Higgins B, Ikeda K, Kanazawa M, Vandergheynst J, Fiehn O, Arita M. 2015. MS-DIAL: Data-independent MS/MS deconvolution for comprehensive metabolome analysis. Nature Methods 12, 523–526.

Tsugawa H, Kind T, Nakabayashi R, Yukihira D, Tanaka W, Cajka T, Saito K, Fiehn O, Arita M. 2016. Hydrogen Rearrangement Rules: Computational MS/MS Fragmentation and Structure Elucidation Using MS-FINDER Software. Analytical Chemistry 88, 7946–7958.

Tun NN, Santa-Catarina C, Begum T, Silveira V, Handro W, Segal Floh EI, Scherer GFE. 2006. Polyamines induce rapid biosynthesis of nitric oxide (NO) in Arabidopsis thaliana seedlings. Plant and Cell Physiology 47, 346–354.

Turko I V., Ballard SA, Francis SH, Corbin JD. 1999. Inhibition of cyclic GMP-binding cyclic GMP-specific phosphodiesterase (type 5) by sildenafil and related compounds. Molecular Pharmacology 56, 124–130.

Ullah S, Kolo Z, Egbichi I, Keyster M, Ludidi N. 2016. Nitric oxide influences glycine betaine content and ascorbate peroxidase activity in maize. South African Journal of Botany 105, 218–225.

Vera-Estrella R, Barkla BJ, Bohnert HJ, Pantoja O. 2004. Novel regulation of aquaporins during osmotic stress. Plant Physiology 135, 2318–2329.

Wong A, Gehring C. 2013. The Arabidopsis thaliana proteome harbors undiscovered multi-domain molecules with functional guanylyl cyclase catalytic centers. Cell Communication and Signaling 11, 48.

Yuan P, Jauregui E, Du L, Tanaka K, Poovaiah BW. 2017. Calcium signatures and signaling events orchestrate plant–microbe interactions. Current Opinion in Plant Biology 38, 173–183.

Yun BW, Feechan A, Yin M, et al. 2011. S-nitrosylation of NADPH oxidase regulates cell death in plant immunity. Nature 478, 264–268.

Zhou J, Jia F, Shao S, Zhang H, Li G, Xia X, Zhou Y, Yu J, Shi K. 2015. Involvement of nitric oxide in the jasmonate-dependent basal defense against root-knot nematode in tomato plants. Frontiers in Plant Science 6, 193.

